# Comparative genomics of smut fungi suggest the ability of meiosis and mating in asexual species of the genus *Pseudozyma* (Ustilaginales)

**DOI:** 10.1101/2023.01.09.523226

**Authors:** Lena Steins, Marco Alexandre Guerreiro, Marine Duhamel, Fei Liu, Qi-Ming Wang, Teun Boekhout, Dominik Begerow

## Abstract

**Background:** The Ustilaginales comprise hundreds of plant-parasitic fungi with a characteristic life cycle that directly links sexual reproduction and parasitism: One of the two mating-type loci codes for a transcription factor that not only facilitates mating, but also initiates the infection process. However, several species within the Ustilaginales have no described parasitic stage and were historically assigned to the genus *Pseudozyma*. Molecular studies have shown that the group is polyphyletic, with members being scattered in various lineages of the Ustilaginales. Together with recent findings of conserved fungal effectors in these non-parasitic species, this raises the question if parasitism has been lost recently and in multiple independent events or if there are hitherto undescribed parasitic stages of these fungi.

**Results:** In this study, we sequenced genomes of five *Pseudozyma* species together with parasitic species from the Ustilaginales to compare their genomic capability to perform two central functions in sexual reproduction: mating and meiosis. While the loss of sexual capability is assumed in certain lineages and asexual species are common in Asco- and Basidiomycota, we were able to successfully annotate functional mating and meiosis genes that are conserved throughout the whole group.

**Conclusion:** Our data suggest that at least the key functions of a sexual lifestyle are maintained in the analyzed genomes, challenging the current understanding of the so-called asexual species with respect to their evolution and ecological role.

## Background

The Ustilaginales are an order of plant-parasitic fungi within the Ustilaginomycotina (Basidiomycota), commonly known as smut fungi of grasses. Although similar plant-parasitic strategies are implemented in other lineages, such as the Urocystidales (Ustilaginomycetes), Tilletiales, or even Doassansiales, Entylomatales (Exobasidiomycetes), and Microbotryaceae (Microbotryomycetes), the sheer number of species in the Ustilaginales express their ecological relevance in certain ecosystems (1). Their adaptations to open, temperate grassland ecosystems include efficient dispersal of diaspores that remain capable of germination for many years (2) and a life cycle with two alternating phases, corresponding to seasonal change. Almost all species in this group share a dimorphic life cycle that comprises an asexual, saprotrophic yeast stage, followed by a filamentous sexual stage that is necessary to parasitize the host (1). Infection is initiated after the mating of two cells of compatible mating-types (i.e., different alleles of mating-type genes). The mating-types are determined by two genetic loci, the pheromone/pheromone receptor locus (PR) and the homeodomain transcription factor locus (HD). The two mating-type loci can be physically linked (bipolar mating) or unlinked (tetrapolar mating) in the genome (3). The pheromone/pheromone-receptor system controls the recognition of compatible mating partners by secretion of small pheromones or mating factors (MFa) that activate the g-protein coupled receptor (PRA) of the partner (4, 5). Generally, one individual produces one or two different pheromone types (species with two (biallelic) or three (triallelic) different alleles in the PR locus) and one receptor type that are unable to activate the mating receptors of cells of the same mating-type, but can activate receptors of strains with different mating-types (i.e., heterothallic), preventing selfing and promoting genetic variability (6). After receptor activation, the cells form conjugation hyphae, proceeding to dikaryotic penetration hyphae to infect the plant (4). This morphological switch is mediated by heterodimers of the homeodomain transcription factor, coded for by the two genes of the HD locus (bEast/bWest). Dimers can only be formed from gene products of two different strains that bear different alleles at the often multiallelic HD locus (7). Several studies indicate that this homeodomain transcription factor not only maintains the dikaryotic growth and controls clamp development as in many Agaricomycotina (Basidiomycota), but it is also a major regulator of the parasitic (teleomorphic) phase in Ustilaginales (8). After the infection of the plant, growth of the parasite is directed towards the site of sporulation, which is either leaf-tissue or, more often, flowers or complete inflorescences. This parasitic phase is characterized by a dikaryotic growth and is finally summarized in the formation of teliospores, which act as resting spores and are dispersed by wind or vectors like insects. Karyogamy and meiosis take place during the germination of the teliospores, resulting in a septated basidia bearing four basidiospores, which can proceed to yeast-like growth of haploid sporidia (anamorphic) (1). The haploid stage can be maintained for a long period of time (1), and initiation of a new infection can only be completed by mating of the sporidia that act as gametes, leading to the formation of the dikaryon and repeating the cycle.

With the advent of molecular phylogenetic tools, it has become clear that some members scattered in the Ustilaginales have only been isolated as yeasts (9). Some of them are found frequently in diverse ecological niches and do not bear sexual or parasitic structures. Thus, the anamorphic genus *Pseudozyma* was used to accommodate the asexual species sharing physiological characteristics of other Ustilaginales (9), while being known for only reproducing asexually and producing compounds relevant for industry and agriculture (10–13). However, the genus has since been found to be polyphyletic and many species have already been reassigned to other genera using molecular taxonomic studies and renamed accordingly (14). In this study, we will use the new names when applicable, but to keep comprehensiveness of the texts, refer to the whole group as ‘*Pseudozyma’*. The reassignment based on molecular phylogenies suggests that asexuality would have evolved multiple times in the evolutionary history of the Ustilaginales (14, 15). This could include the loss of mating and meiosis machinery that are only required in sexual reproduction, as well as the adaptation from a nutritional diverse metabolism (16) that is characterized by switching between saprobic and parasitic stages to purely saprobic growth.

As proposed by Begerow et al. (9), some *Pseudozyma* species have been found to represent the anamorphic stage of described parasitic species, a teleomorph for *Pseudozyma antarctica* (*Moesziomyces antarcticus*) has been described recently (17), as well as cases of probable conspecificity in the *Moesziomyces* genus (15). Adding on that, Sharma et al. (18) have annotated fast-evolving parasitic effector sets in several *Pseudozyma* species *in silico* and provided experimental evidence that the conserved effector *Pep1* that is known from the model organism *Ustilago maydis* is present and functional in *Pseudozyma*. However, it remains unclear if this is true for all *Pseudozyma* species and whether their parasitic stages have been missed until now, or if there are truly asexual species which have lost their capacity for mating and meiosis. There are several scenarios which would support such hypotheses, such as the extinction of the host or the lack of a mating partner with the compatible mating genes. In addition, symptomless colonization could explain why the sexual, plant-parasitic stages have not been observed so far.

The presence of effector genes relevant for parasitism in *Pseudozyma* raises the question if they share the genomic makeup of sexual Ustilaginales species to proceed through the typical life cycle of this group. To answer this question, we annotated the core genes for two central sexuality-related features in the life cycle of the fungi: mating and meiosis. This includes both mating loci (PR and HD), as well as 20 core meiosis genes described for Ustilaginomycotina (19–21). We tested the functionality and conservation of the annotated genes *in silico*. We subsequently inferred possible mating systems (bi-or triallelic, bi- or tetrapolar) in *Pseudozyma* and other related Ustilaginales species and assessed conservation of meiosis-related genes to hypothesize possible sexual reproduction in *Pseudozyma*. We found that the *Pseudozyma* species all display a functional genetic makeup for sexual reproduction, challenging their anamorphic species assignment.

## Results

### Genome sequencing of Ustilaginales

In this study, we sequenced and assembled 11 genomes from Ustilaginales strains (belonging to 11 different species from five genera including *Pseudozyma*). Genome statistics were computed to assess the genome structure and quality of newly sequenced and published genomes for comparison (Table 1). The genome size of the newly sequenced genomes ranged from 13.8 Mb in *Farysia itapuensis* (*Farysizyma itapuensis*) to 24.8 Mb in *Sporisorium sorghi*. These genomes extended the range of the published genomes incorporated in this study, comprising sizes from 17.3 to 20.8 Mb. The quality of the assembly of the Illumina reads varied between 71 scaffolds in *Farysia itapuensis* and 6454 scaffolds in *Ustilago tritici*, while published genomes were assembled in 22 to 12840 scaffolds, respectively. N50 values of new genomes ranged between 18.8 kb and 473.1 kb. However, the assemblies contained less gaps than some of the already published genomes. This might have led to a higher scaffold count but reduces the risk of misassembly by scaffolding or problems due to masking.

**Table 1:** Genome and annotation statistics. Genomes of species formerly classified in the *Pseudozyma* genus (asexual) are marked in bold.

| Species name | Strain | Size (Mb) | Scaffolds | N50 | Gaps (n) | GC content | Genes | Introns/gene | Genes/Mb | BUSCO | Source | Accession |
| --- | --- | --- | --- | --- | --- | --- | --- | --- | --- | --- | --- | --- |
| *Pseudozyma/Anthracocystis flocculosa* | CBS 167.88.2 | 18.7 | 83 | 391904 | 8 | 57.15% | 7417 | 0.57 | 397 | 99.1% | This study | GCA_023212635.2 |
| *Farysia itapuensis* | CBS 10428 | 13.8 | 71 | 473075 | 14 | 54.4% | 5791 | 0.65 | 421 | 98.2% | This study | GCA_023212645.1 |
| *Kalmanozyma brasiliensis* | GHG001 | 17.3 | 45 | 720612 | 117 | 58.1% | 6394 | 0.61 | 370 | 99% | (66) | GCA_000497045.1 |
| *Moesziomyces antarcticus* | JCM 10317 | 18.1 | 197 | 701213 | 97 | 60.9% | 6727 | 0.83 | 372 | 98.3% | (67) | GCA_000747765.1 |
| *Moesziomyces aphidis* | DSM 70725 | 17.9 | 42 | 713260 | 2089 | 61.2% | 6623 | 0.88 | 370 | 95.9% | (68) | GCA_000517465.1 |
| *Moesziomyces parantarcticus* | CBS 10005 | 18.2 | 198 | 258071 | 0 | 60.2% | 6727 | 0.8 | 369 | 98.7% | This study | GCA_023212605.1 |
| *Pseudozyma hubeiensis* | SY62 | 18.4 | 74 | 445580 | 86 | 56.5% | 6465 | 0.6 | 351 | 97.8% | (11) | GCA_000403515.1 |
| *Pseudozyma pruni* | CBS 10937 | 17.9 | 114 | 319176 | 15 | 57.8% | 6373 | 0.67 | 357 | 99.1% | This study | GCA_023212685.1 |
| *Pseudozyma tsukubaensis* | NBRC 1940 | 23.8 | 12840 | 319176 | 0 | 53.4% | 8889 | 0.56 | 373 | 98.8% | (55) | GCA_001736125.1 |
| *Pseudozyma thailandica* | CBS 10006 | 19.7 | 1899 | 461922 | 21 | 53.6% | 6570 | 0.56 | 334 | 98.9% | This study | GCA_023212835.1 |
| *Sporisorium graminicola* | CBS 10092 | 19.6 | 22 | 966477 | 4 | 56.8% | 6619 | 0.68 | 338 | 99.0% | (12) | GCA_005498985.1 |
| *Sporisorium reilianum* f. sp. *reilianum* | SRS1_H2-8 | 18.5 | 23 | 782463 | 944 | 59.7% | 6591 | 0.69 | 356 | 98.5% | (69) | GCA_900162835.1 |
| *Sporisorium scitamineum* | CBS 131463 | 19.6 | 371 | 94037 | 47 | 55.0% | 6604 | 0.7 | 338 | 98.4% | This study | GCA_023212615.1 |
| *Sporisorium sorghi* | CBS 104.17 | 24.8 | 782 | 67233 | 192 | 55.5% | 8577 | 0.88 | 345 | 98.5% | This study | GCA_023212695.1 |
| *Testicularia cyperi* | MCA 3645 | 20.8 | 95 | 1344767 | 157 | 56.4% | 6201 | 0.7 | 298 | 97.5% | (70) | GCA_003144125.1 |
| *Tranzscheliella williamsii* | CBS 131475 | 18.5 | 241 | 219023 | 6 | 57.9% | 6403 | 0.93 | 347 | 98.3% | This study | GCA_023212725.1 |
| *Urocystis primulicola* | RUB 030670 | 13.8 | 26 | 619582 | 0 | 54.1% | 5355 | 1.7 | 388 | 97.6% | JGI: mycocosm | PRJNA456443 |
| *Ustilago hordei* | CBS 131470 | 21.3 | 2359 | 18849 | 35 | 51.5% | 6323 | 0.81 | 297 | 97.9% | This study | GCA_023212755.1 |
| *Ustilago maydis* | FBA (CBS 132774) | 19.5 | 158 | 292465 | 25 | 54.0% | 6332 | 0.51 | 325 | 98.9% | This study | GCA_023212765.1 |
| *Ustilago tritici* | W66970 (CBS 669.70) | 22.0 | 6454 | 74669 | 60 | 52.5% | 8228 | 0.72 | 374 | 99.0% | This study | GCA_023213265.1 |

The BUSCO genome completeness assessment resulted in completeness values between 97.8% (*Ustilago hordei*) and 99.1% (*Pseudozyma pruni*), published genomes were between 95.9% (*Moesziomyces aphidis*) and 99% (*Sporisorium graminicola*) complete.

*De novo* gene prediction resulted in between 5,355 genes for the outgroup species *Urocystis primulicola* and 8,889 genes for *Pseudozyma tsukubaensis*. Resulting gene densities ranged between 297 genes/Mb in *U. hordei* and 421 genes/Mb in *Farysia itapuensis*, the smallest genome in the study. The lowest number of introns per gene was predicted in *Ustilago maydis* (0.51), the highest in *Urocystis primulicola* (1.7). All Ustilaginales genomes in this study contained less than one intron per gene (Table 1).

Analyzing the predicted genes with OrthoFinder resulted in 2,337 single-copy orthologous genes that were aligned for calculating a phylogenetic tree with *Urocystis primulicola* as outgroup (Fig. 1). This shows that *Pseudozyma* is a polyphyletic genus, validating reassignments of species to other genera, e.g., *Anthracocystis, Sporisorium*, and *Moesziomyces* (14).

**Figure 1:**
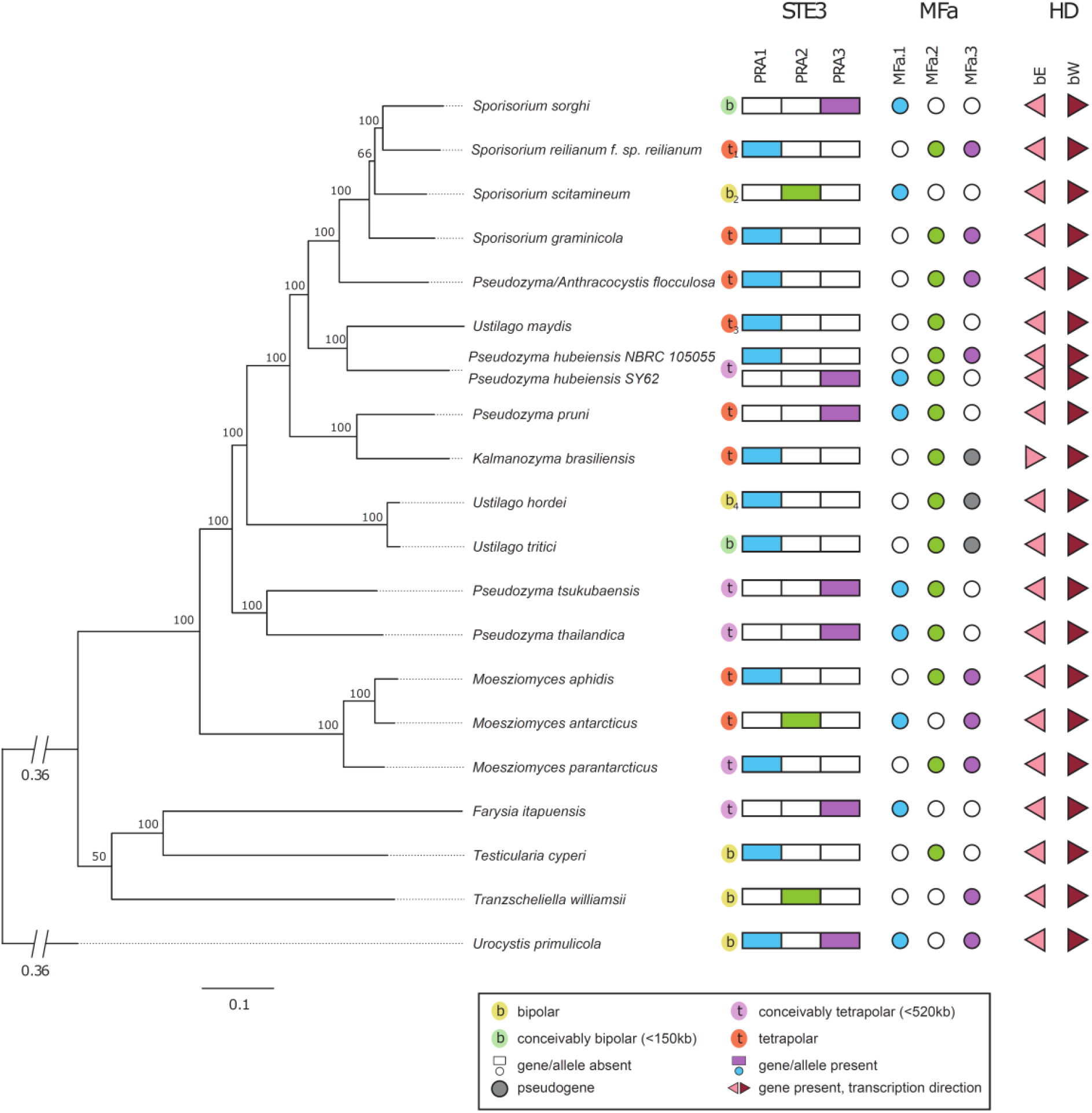
Phylogenomic tree of *Ustilaginales* and annotation of mating genes. The phylogeny was calculated based on 2,337 single copy orthologous genes. *Urocystis primulicola (Urocystidales)* was specified as outgroup. *Pseudozyma hubeiensis* (strain NBRC 105055) was not included in the calculation of the tree but added to the figure to visualize the mating genes. ^1^ Tetrapolar (24), ^2^ Bipolar (22), ^3^ Tetrapolar (2), ^4^ Bipolar (23)

### Genomes of *Pseudozyma* show functional mating-type loci

Manual annotation by multiple sequence alignments revealed that all genomes in the study contained putative functional genes coding for one pheromone receptor and one or two pheromones, as well as for both homeodomain transcription factor subunits (bEast/bWest) (Fig. 1).

Almost all *Pseudozyma* genomes contained one pheromone receptor allele and two different pheromones at the a-locus (Fig. 1). The only exception was *Kalmanozyma brasiliensis* that only contained one functional pheromone copy, with the other one lacking the stop codon. Some sexual species (*Ustilago hordei, Ustilago tritici*, and *Ustilago maydis*), also showed a second pheromone gene that seemed to be pseudogenized (i.e., strongly resembling the overall sequence of the respective pheromone, but lacking the stop codon). Other species, such as *Sporisorium reilianum, Sporisorium sorghi, Sporisorium scitamineum*, and *Testicularia cyperi*, maintained only one pheromone without showing evidence for a pseudogenized pheromone. For *Pseudozyma hubeiensis*, strain SY62 showed an a_3_ mating-type, carrying pheromones for a_1_ and a_2_, and strain NBRC 105055 an a_1_ mating-type with pheromones for a_2_ and a_3_.

We were able to resolve the allele of each receptor and pheromone gene based on phylogenetic analyses (Supplementary Fig. 1 and 2), and therefore identified the mating-type for each strain based on the receptor. For all genomes, we identified that the pheromone receptor coding genes belonged to one allele, while the pheromone genes only were compatible to other alleles.

The HD locus, including functional domains of the genes (Supplementary Table 1), could also be annotated for all species and was structured in the same way in almost all genomes, with the genes being adjacent and divergently transcribed (Fig. 1). In *Kalmanozyma brasiliensis*, the structure of the HD locus deviated, as both genes were located on the same strand. The two sequenced *Pseudozyma hubeiensis* strains showed differences in the HD locus genes that imply different alleles of the bEast and bWest genes.

Additionally, we assessed bipolarity and tetrapolarity in the species (Fig. 1). Both mating-type loci were located on the same scaffold in *Testicularia cyperi, Tranzscheliella williamsii*, and *Urocystis primulicola* (genome was sequenced from a culture containing both mating-types) with a genomic distance of less than 520 kb. For non-chromosomal assemblies, the determination of tetra- and bipolarity remains challenging if flanking regions of the mating-type loci were not correctly assembled and different scaffolds could be part of the same chromosome. Addressing this, telomeres were predicted for the scaffolds containing both mating-type loci (for comparative purposes, we included chromosomal assemblies). *Moesziomyces antarcticus, Sporisorium graminicola*, and *Kalmanozyma brasiliensis* showed telomeres at one or both scaffolds containing mating-type loci on both ends (Supplementary Table 2), establishing the tetrapolarity of these species. For *Pseudozyma hubeiensis* and *Pseudozyma/Anthracocystis flocculosa*, only one telomere could be annotated on the distal end of one mating scaffold, not providing information exceeding the minimal total flanking regions. Adding on this, the PR locus in *Pseudozyma/Anthracocystis flocculosa* was annotated at the ends of two different scaffolds, split between the pheromone receptor and the pheromones.

*Sporisorium sorghi* and *Ustilago tritici* were assessed to be bipolar, as the loci were located on different scaffolds with minimal total assembled flanking regions smaller than 150 kb. Although *Ustilago hordei* and *Sporisorium scitamineum* were previously identified as bipolar species (22, 23), we could not retrieve this result in our study due to assembly quality. *Farysia itapuensis, Moesziomyces parantarcticus, Pseudozyma hubeiensis, Pseudozyma thailandica*, and *Pseudozyma tsukubaensis* could be either pseudobipolar (mating-type loci located on the same chromosome but inherited independently) or tetrapolar, as their mating-type loci were located on two different scaffolds and the minimal total assembled flanking regions were between 150 kb and 520 kb. All other species are tetrapolar according to definition (located on two different scaffolds and more than 520 kb distance minimal total assembled flanking regions or located on two different scaffolds and tetrapolar according to previous studies (2, 24). Many of the species in our study that only had one pheromone or a pseudogenized pheromone are bipolar. However, *Farysia itapuensis* could be tetrapolar, as well as *Kalmanozyma brasiliensis. Ustilago maydis*, which also contained a pseudogenized pheromone, is a known tetrapolar species (25).

### *Pseudozyma* species have the same genomic capability of meiosis as sexual species of Ustilaginales

Most core meiosis genes from *Ustilago maydis* were detected and identified as potentially functional in all tested species using manual annotation with BLAST and multiple sequence alignments (Fig. 2). Functional gene models were lacking in *Moesziomyces aphidis* (SMC1, SMC2) due to gaps in the assembly within the potential gene models. However, the parts of the gene that flanked the gap could be aligned with the reference gene and coding sequence of *Ustilago maydis* without premature stop codons. Treating the gaps as introns, a functional gene model could be produced. In *Ustilago hordei* and *Sporisorium sorghi*, MER3 and SPO11, respectively, were located at the ends of scaffolds and could therefore not be modeled completely, despite being detected in the genomes. In the outgroup species *Urocystis primulicola*, SMC6 and SPO11 could not be annotated, as even extensive tblastn searches did not produce an alignment specific to these genes. In summary, all *Pseudozyma* species besides *Moesziomyces aphidis* produced reliable gene models for core meiosis genes, resembling sexual species in the *Sporisorium* and *Ustilago* genera, and other, less closely related sexual species, such as *Testicularia cyperi* and *Tranzscheliella williamsii*. Alignments with the meiosis specific genes MER3, MSH4, REC8, and SPO11 produced functional gene models in all *Pseudozyma* and most sexual species in Ustilaginales. Meiosis gene loci were syntenic between the analyzed species corresponding to their phylogenetic relations, showing little rearrangements between closely related species. No rearrangements could be specifically linked to species that have been described to be asexual (Supplementary Fig. 3).

**Figure 2:**
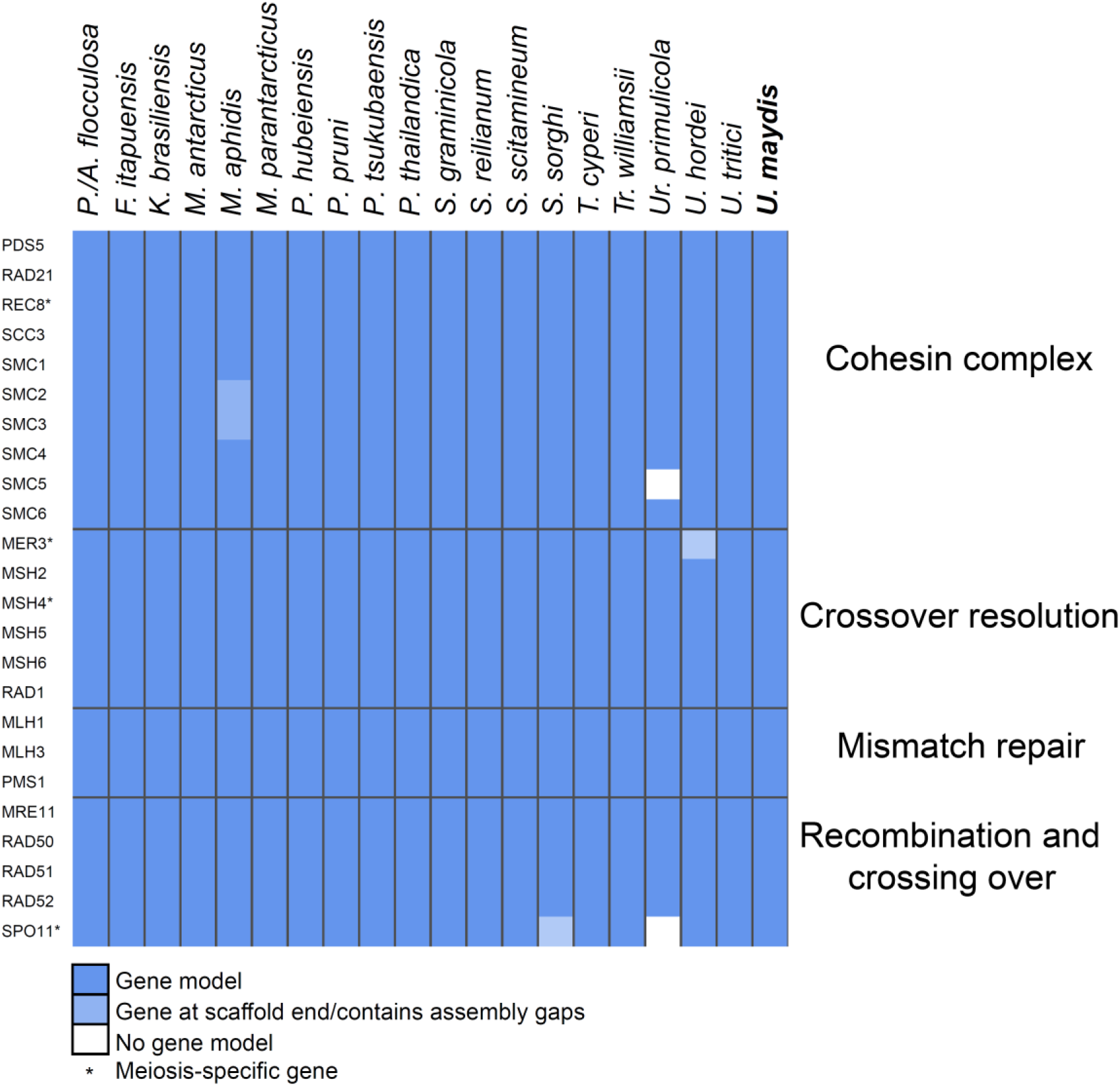
Heatmap showing presence of functional core meiosis genes from *Ustilago maydis* in other species. Curated gene models were created by multiple sequence alignments.

To further assess possible functionality of meiosis and mating gene models, a HMMER protein domain search was performed on the translated coding sequences of the genes using the Pfam database. Compared to *Ustilago maydis* data from Pfam and InterPro as a reference, the gene models produced in this study show all specific functional domains of the respective reference genes (Supplementary Table 1). The location of the domains in the genes was syntenic to their homologs in *Ustilago maydis* in all tested strains. However, the SMC3 gene model in *Moesziomyces aphidis* and the SMC4 gene model in *Urocystis primulicola* each showed a shortened SMC_N-domain. Despite these, there were no length or location irregularities in the functional domains of the annotated genes in the *Pseudozyma* genomes.

We then investigated the relaxation of selection on genes involved in or specific to meiosis in *Pseudozyma* using RELAX. The program compares selection in a test group (*Pseudozyma*) to a reference group (sexually reproducing species) and computes a coefficient *k* which suggest relaxation (*k*<1), no change (*k*=1), or intensification (*k*>1) of selection in the test group (*Pseudozyma*). We detected intensified selection on 13 genes that are not specific for but involved in meiosis (Table 2). These genes are known to be involved in processes like mitosis and DNA repair additionally to meiosis (Table 3). Especially strongly intensified selection could be detected for RAD51, that is involved in DNA repair and recombination. Nine other genes showed no difference in selective pressure compared to sexual species, including MSH4 and REC8, which are meiosis-specific genes. Slightly relaxed selection (k=0.91) was observed for MER3, a meiosis specific gene. SPO11, which is essential for meiosis by creating double strand breaks during crossing-over (26), showed more strongly relaxed selection (k=0.59). However, SPO11 could not be detected in the sexual outgroup species *Urocystis primulicola*.

**Table 2:** Selection on meiosis genes in *Pseudozyma* compared to sexual Ustilaginales. The RELAX analysis indicates relaxed and intensified selection on meiosis genes. Relaxed selection was detected for MER3 and SPO11, while selection on the other genes was either intensified or no differences were detected (no k-value provided). Meiosis-specific genes are marked with an asterisk.

| Gene | p | K | Selection |
| --- | --- | --- | --- |
| MER3* | 0.0283 | 0.91 | Relaxed |
| MLH1 | 0.05 | 1.74 | Intensified |
| MLH3 | 0.331 |  | - |
| MRE11 | 0.05 | 3.01 | Intensified |
| MSH2 | 0.05 | 1.46 | Intensified |
| MSH4* | 0.15 |  | - |
| MSH5 | 0.397 |  | - |
| MSH6 | 0.05 | 1.82 | Intensified |
| PDS5 | 0.05 | 1.36 | Intensified |
| PMS1 | 0.573 |  | - |
| RAD1 | 0.0001 | 1.42 | Intensified |
| RAD21 | 0.05 | 1.33 | Intensified |
| RAD50 | 0.115 |  | - |
| RAD51 | 0.0029 | 11.09 | Intensified |
| RAD52 | 0.091 |  | - |
| REC8* | 0.858 |  | - |
| SCC3 | 0.05 | 1.68 | Intensified |
| SMC1 | 0.0008 | 1.2 | Intensified |
| SMC2 | 0.0139 | 1.19 | Intensified |
| SMC3 | 0.772 |  | - |
| SMC4 | 0.05 | 1.87 | Intensified |
| SMC5 | 0.05 | 1.43 | Intensified |
| SMC6 | 0.204 |  | - |
| SPO11* | 0.0124 | 0.59 | Relaxed |

**Table 3:** Core meiosis genes from *Ustilago maydis* and their function during meiosis. Functions listed were obtained from Gene Ontology (56, 57) accessed via UniProt 11/2020 (58). Meiosis specific genes are marked with an asterisk.

| Gene name | *U. maydis* locus tag | Gene product function (GO) | NCBI Gene ID |
| --- | --- | --- | --- |
| MER3* | UMAG_11008 | meiosis specific DNA helicase | 23566946 |
| MLH1 | UMAG_05208 | protein required for mismatch repair in mitosis and meiosis | 23565158 |
| MLH3 | UMAG_03481 | protein involved in DNA mismatch repair and meiotic recombination | 23563923 |
| MRE11 | UMAG_04704 | double-strand break repair protein | 23564799 |
| MSH2 | UMAG_03025 | protein that binds to DNA mismatches | 23563613 |
| MSH4* | UMAG_05846 | protein involved in meiotic recombination | 23565623 |
| MSH5 | UMAG_12155 | DNA mismatch repair protein | 23567913 |
| MSH6 | UMAG_11009 | protein required for mismatch repair in mitosis and meiosis | 23566947 |
| PDS5 | UMAG_03740 | mitotic and meiotic cohesin maintenance factor | 23564110 |
| PMS1 | UMAG_01932 | ATP-binding protein for mismatch repair | 23562798 |
| RAD1 | UMAG_10396 | single-stranded DNA endonuclease | 23566436 |
| RAD21 | UMAG_02591 | mitotic cohesin complex, double-strand-break repair protein | 23563304 |
| RAD50 | UMAG_01085 | DNA repair protein | 23562202 |
| RAD51 | UMAG_03290 | DNA repair protein, strand exchange protein | 23563788 |
| RAD52 | UMAG_04989 | DNA repair and recombination protein | 23565003 |
| REC8* | UMAG_00172 | meiotic cohesin complex subunit | 23561549 |
| SCC3 | UMAG_02053 | sister chromatid cohesion protein | 23562894 |
| SMC1 | UMAG_12218 | subunit of the cohesin complex | 23567971 |
| SMC2 | UMAG_05835 | subunit of the cohesin complex | 23565615 |
| SMC3 | UMAG_04389 | subunit of the cohesin complex | 23564588 |
| SMC4 | UMAG_12209 | subunit of the cohesin complex | 23567962 |
| SMC5 | UMAG_11074 | subunit of the SMC5-SMC6 complex | 23567003 |
| SMC6 | UMAG_00739 | subunit of the SMC5-SMC6 complex | 23561954 |
| SPO11* | UMAG_10420 | meiosis-specific topoisomerase | 23566457 |

## Discussion

To assess the capability of mating and meiosis of members of the asexual (former) genus *Pseudozyma*, we used available and newly sequenced genomes of members of the order Ustilaginales. These genome sequences enabled us to support the recent, marker-based phylogenies that were used to position *Pseudozyma* species in other genera (14) with a multigene phylogeny. The overall topology of Ustilaginales as proposed by Begerow et al. (1) or Wang et al. (14) received higher support through our phylogenomic approach (Fig. 1), and most clades were well-resolved. The whole genome data revealed no tendency for *Pseudozyma* species to harbor smaller or bigger genomes compared to their parasitic relatives (Table 1). The number of predicted genes and low amount of introns per gene was also typical for Ustilaginales and does not differentiate *Pseudozyma* species from plant-parasitic species (27, 23). Furthermore, members of *Pseudozyma* did not show changes in their genetic makeup regarding meiosis and mating, two important keys to the parasitic life cycle that is typical for smut fungi.

### Mating

We were able to detect the core mating genes of both the PR locus and the HD locus in the *Pseudozyma* genomes and assess the functionality of the genes by comparing them to the mating genes of known parasites (Fig. 1). Our findings support previous detections of mating genes in some *Pseudozyma* strains that have been published (10, 28) with the information from newly sequenced *Pseudozyma* species (Fig. 1), assignment of mating-types (Fig. 1, Supplementary Fig. 1 and 2) as well as an assessment of possible functionality (Table 2, Supplementary Table 1, Supplementary Fig. 3).

However, mating genes have previously been detected in representatives of the putatively asexual human pathogenic species of *Malassezia* (Malasseziales) (19), which could imply that mating related genes are maintained despite the potential loss of sexual reproduction. Assuming that members of *Pseudozyma* are able to mate, we showed that they would be self-sterile (heterothallic), containing either one or two pheromones compatible with other mating-types but not their own receptor, as all other known members of Ustilaginaceae (29). In the two *Pseudozyma hubeiensis* strains, we were able to identify two different alleles for the pheromone receptor gene, inter-strain compatible pheromones, and divergent HD genes, which would allow those two strains to conjugate, enter, and maintain filamentous growth. This strain pair could therefore be a candidate for culture-based mating assays to show the capability of sexual reproduction independently of the unknown potential host plant.

In *Kalmanozyma brasiliensis*, the genes at the HD locus were organized adjacently and in the same orientation rather than being divergently transcribed, which is a deviation of the usual organization in Basidiomycota (29) and in all other species in this study. Nevertheless, both genes were annotated with full functional domains and are therefore potentially functional in sexual reproduction, i.e., potential filamentous growth that is maintained by the HD dimers (8).

The presence of two different pheromones in most analyzed *Pseudozyma* genomes hints to the existence of three different pheromone receptor alleles in these species, as found in many other species of Ustilaginaceae (6). The possible pseudogenized pheromone in *Kalmanozyma brasiliensis*, as well as in some sexual species of Ustilaginales, is congruent to the situation in *Ustilago maydis* that displays a tetrapolar and biallelic mating system with a pseudogenized pheromone (25, 5). Non-chromosomal assemblies of most of our data do not allow a thorough analysis of bipolarity and tetrapolarity, which is best shown in the examples of *Farysia itapuensis* and *Pseudozyma/Anthracocystis flocculosa*, where even the PR locus seems to be incorrectly assembled and split on the ends of two different scaffolds. This displays the need for more long-read sequencing data in the Ustilaginales order to study mating system evolution, also taking into account the accumulation of transposable elements that is expected for bipolar species evolving from the tetrapolar ancestral state (22, 29), as recently shown in the anther smut genus *Microbotryum* (Microbotryales) (30). Nevertheless, our data support the plesiomorphic situation with three alleles of the PR locus as previously suggested (6) and hint towards a situation of scattered bipolarity and tetrapolarity. This would mirror the situation in the lineage of *Microbotryum*, in which multiple independent events are responsible for bipolar mating systems in most species (30, 31).

### Meiosis

We were able to detect and annotate all core meiosis gene as defined for *Ustilago maydis* (19) in the genomes from *Pseudozyma* strains (Fig. 2) and showed that most of them displayed no relaxed selection in the examined genomes (Table 2). In addition, all core meiotic genes have shown syntenic locus organization in closely related species throughout all meiotic functions (Supplementary Fig. 3) and maintained functional protein domains (Supplementary Table 1). The maintenance of meiosis genes strongly suggests the relevance of meiosis in *Pseudozyma*, as it is likely that genes that are not utilized in an organism will be lost or pseudogenized eventually (32). Another possibility would be a neofunctionalization, which would include increased or delayed mutation rates in these genes. If this case, we would expect a gain or loss of functional domains in the genes and a striking difference in gene conservation for *Pseudozyma* species compared to sexual species, as well as genomic rearrangements at the meiosis gene loci that cannot purely be explained by species borders. The comparison of selection strength on the meiosis-specific genes SPO11 and MER3 showed a relaxation of selection in the *Pseudozyma* group (Table 2) compared to the other species. SPO11 is widely considered to be essential for meiosis (26), although it has been lost in a lineage of amoeba that performs meiosis (33, 34) and could not be successfully annotated in the sexually reproducing outgroup species *Urocystis primulicola*. This could be a sign that it is non-essential in smut fungi, but more likely the sequence conservation is less striking than in other meiosis-related genes and the gene could not be identified by our strategy. Meiosis-specific genes REC8 and MSH4 and other meiosis-related genes showed no change in selection in *Pseudozyma* compared to other species, leading to the conclusion that the biological function of meiosis is conserved over all analyzed Ustilaginales, as it is in eukaryotes in general (20, 35, 36). In contrast, *Malassezia* species lack some relevant core genes like MSH5, or meiosis-specific genes MSH4 and MER3, while *Pseudozyma* species typically retained all 20 core meiosis genes found in *Ustilago maydis* (19, 20, 36). Thus, we conclude that the analyzed species of *Pseudozyma* seem to be capable of meiosis, as members of the genus *Malassezia* have probably lost a considerable part of core meiosis genes, while still showing some evidence for recombination and hybridization (19, 37). If there was a loss of function in meiosis genes in *Pseudozyma*, we hypothesize it to be on protein-structure level and probably very recent, as the functional domains of the genes are conserved and seem functional. Additionally, such a change would have occurred multiple times throughout evolution while maintaining syntenic locus organization without any hints for increased genomic rearrangement activity around the core meiosis genes.

### Conclusion

The genetic makeup of *Pseudozyma* species in mating and meiosis loci suggests not only that these species are able to sexually reproduce, but, given the close linkage of sexual reproduction and parasitism in Ustilaginales, could colonize plants causing the smut syndrome. This is supported by core effector genes that could be annotated for some of the species included in this study, and by the functional conservation of the *Pep1* effector that is responsible for establishing interaction zones with the host plant in *Ustilago maydis* (18). It could be possible that most species labeled as *Pseudozyma* can be assigned as the yeast stage of known parasites utilizing genomic and other molecular data in the future (15), or new collection trips may reveal their parasitic stage of so far unknown host plants (17, 15). A different scenario would be that these smuts cause little to no symptoms when colonizing the host plant, maybe as endophytes. That can make it difficult to sample the sexual stage, although some *Pseudozyma* species can be found in association with plants (13). Another possibility are parasitic species with a strongly dominating yeast stage, as previously suspected for the genus *Moesziomyces* (15, 38). This would, much like the loss of sexual stages, have evolved multiple times through-out the phylogenetic tree of the Ustilaginales, but it might be favorable for the fungi to avoid immune responses of the host, while the complete loss of sexual reproduction is characterized by a loss of genetic variability which would probably lead to extinction in the long run (39).

## Material and Methods

### Sequenced strains and incorporated published genomes

For this study, we sequenced 11 genomes from Ustilaginales species (Table 1). For each species, a haploid strain was selected. All strains were obtained from existing pure cultures deposited at the fungal culture collection of the Westerdijk Fungal Biodiversity Institute (CBS). Five of these strains belong to species in the former *Pseudozyma* genus (*Pseudozyma/Antracocystis flocculosa, Moesziomyces parantarcticus, Pseudozyma pruni, Pseudozyma tsukubaensis, Pseudozyma thailandica*). We sequenced additional genomes of *Ustilago hordei, Ustilago tritici*, and *Ustilago maydis* for comparable data, as well as strains *of Sporisorium sorghi, Tranzscheliella williamsii*, and *Farysia itapuensis* (*Farysizyma itapuensis*). For further comparison of sexual and asexual species, we incorporated published genomes of former *Pseudozyma* species (*Kalmanozyma brasiliensis, Moesziomyces aphidis, Moesziomyces antarcticus, Pseudozyma hubeiensis, Sporisorium graminicola*) and sexual species (*Sporisorium reilianum, Sporisorium scitamineum, Testicularia cyperi*) in the analysis, and used a sexually reproducing representative from the sister group Urocystidales (*Urocystis primulicola*, strain sequences from a culture containing cells from both mating-types) as outgroup. *Moesziomyces aphidis* and *Moesziomyces antarcticus* were classified in the *Pseudozyma* group despite the possibility of conspecificity with sexual morphs and the recent discovery of a teleomorph, respectively, to examine them for possible signs of reduced sexual reproduction as proposed for the genus (38).

### DNA extraction

Ustilaginales strains were incubated in liquid medium in a 250 mL Erlenmeyer flask at 25°C. After 48-72 h, we retrieved the cells, then snap froze them in liquid nitrogen. We extracted genomic DNA using the CTAB method (40, 41): We used a sterile mortar and pestle to disrupt the frozen cells, then rapidly transferred the resulting fine powder to a 1.5 mL microfuge tube and added DNA lysis buffer (100 mM Tris (pH 8.0), 20 mM EDTA, 2% CTAB, 1.2 M NaCl, and 0.1% β-mercaptoethanol) preheated in a 60 °C water bath. We vortexed the sample thoroughly and incubated it at 60°C for 1 h. The mixture was cooled to room temperature, then admixed with an equal volume of phenol: chloroform: isoamyl alcohol (25:24:1, v/v/v), and centrifuged at 10,000 rpm for 15 min. We transferred the supernatant to a new microtube, and added an equal volume of cold absolute isopropanol for precipitating total DNA at -20°C for 20 min; we then centrifuged the mixture at 10,000 rpm for 10 min. We washed t pellet twice with 75% ethanol and centrifuged it at 10,000 rpm at 4°C for 10 min. We discarded the supernatant, and dried the pellet at RT. We resuspended the DNA pellet in 500 μl high salt TE (10 mM Tris (pH8.0), 2 mM EDTA, 1 M NaCl) with 2.5 μl RNase A at 60°C for 1 h. We added an equal volume of chloroform:isoamyl alcohol (24:1, v/v) and centrifuged the sample at 10,000 rpm for 15 min. We transferred the supernatant to a new microtube and added 2 volumes of precooled ethanol. We inverted and then centrifuged the tube at 10,000 rpm for 10 min to pellet the DNA. We removed the supernatant and dried the pellet at RT, before resuspending it in 100 μL TE buffer. We stored the extracts at −4 °C for immediate use or at −20 °C for long-term storage.

### Genome sequencing and assembly

All sequencing libraries were constructed using the Illumina TruSeq library kit. Paired-end sequencing (150 bp/read) was performed using an Illumina NovaSeq 6000 sequencer.

We trimmed adapters of the raw reads and filtered low-quality sequences using fastp v0.19.5 (42) with default options. We then *de novo* assembled the genome sequences then with SPAdes v3.13.1 (43) using the “– careful” and “– cov-cutoff auto” parameters.

We applied BUSCO version 5.1.3 (44) to assess genome completeness using the included Basidiomycota lineage data (basicdiomycota_odb10) as a reference.

### Phylogenetic tree reconstruction

We predicted genes in assembled genomes using AUGUSTUS version 3.3.2 (45) pre-trained for *Ustilago maydis* and obtained the corresponding translated sequences as fasta files with an integrated script. We identified orthologous genes using OrthoFinder version 2.3.1 (46) and individually aligned the resulting 2,337 single copy orthologuous sequences present in all species using MAFFT version 7.427 (47). We concatenated the alignments with the seqkit package (48) and calculated a multigene phylogeny in RaxML version 8 (PROTGAMMAWAG substitution model, rapid Bootstrap analysis, 500 runs (49)), specifying *Urocystis primulicola* as outgroup.

### Manual meiosis and mating gene annotation

To identify core meiosis and mating genes, we retrieved reference genes from *Ustilago maydis* (meiosis genes, Table 3) and other species (mating genes, Supplementary Table 3) from NCBI database as whole gene sequences, coding sequences, and amino acid sequences. We applied the tblastn algorithm version 2.4.0 (50, 51) to identify these reference genes in the genomes by curating the alignments manually (pre-filtered for e≤ 0.0001). We performed multiple sequence alignments of the candidate loci including their immediate flanking regions using MAFFT version 7.427 (47) with the full gene and coding nucleotide sequences of the respective reference genes. Based on these alignments, we performed several translation steps and multiple alignment steps in MEGA version 5 (52), utilizing the integrated MUSCLE algorithm (53) to manually model start and stop codons as well as introns and exons based on the reference genes’ reading frames and local gene prediction with AUGUSTUS online tool pre-trained for *Ustilago maydis* (54).

We extended the identification of the mating genes to a second sequenced strain of *Pseudozyma hubeiensis* (strain NBRC 105055, Accession GCA_001736105.1, 55) to assess the possibility of two different mating-types for this species.

### Assignment of mating-types and systems

After modeling the pheromone receptor and pheromone genes, the sequences into amino acid sequences. For the PRA and MFa genes, respectively, we aligned the modeled genes together with published, mating-type assigned sequences from NCBI database (Supplementary Table 3) using MAFFT (47) and calculated gene trees using RAxML v8 (PROTGAMMAWAG substitution model, rapid Bootstrap analysis, 1000 runs (49)). From the location of the modeled genes in the tree clusters, we inferred the alleles of the genes. For a better visualization, we recalculated the trees using only gene models from the studied genomes after assignment of mating-types.

We assessed the PR and HD loci linkage by i) scanning the scaffolds containing mating genes for telomeric regions and ii) calculating the minimal total assembled flanking regions between the loci if they were located on different scaffolds, assuming that the scaffolds could be connected to form a chromosome (in absence of telomeric repeat sequences). We predicted telomeres by scanning the output of a Tandem Repeats Finder version 4.10.0 (59) analysis (recommended parameters) for the telomeric repeat pattern known for *Ustilago maydis* TTAGGG (60) at the beginning or end of scaffolds.

We assumed non-linked mating-type loci (tetrapolar) when the minimal total assembled flanking regions were longer than 520 kb (biggest known distance of mating-type loci in a bipolar Ustilaginomycete (*Ustilago hordei* (22, 29), excluding flanking regions that contained a telomeric sequence) or one or both scaffolds containing the mating-type loci were assembled with both telomeres. We assumed a tendency for possible tetrapolarity for regions between 150 kb and 520 kb, and a tendency for possible bipolarity for regions shorter than 150 kb (distance of mating-type loci in bipolar species like *Malassezia sympodialis* (19)). We assigned bipolarity when the loci were located on the same scaffold with a distance of less than 520 kb (22, 29).

### Conservation of meiosis genes

To assess the functionality of meiosis and mating genes, we detected functional domains using the standalone Pfam database v 33.0, (61) using HMMER version 3.3 (62). We analyzed the resulting annotated protein domains in comparison to the respective reference genes from *Ustilago maydis* in the Interpro (63) and Pfam (61) online databases. We checked functional protein domains for full length and relative location within the sequence.

We established conservation analyses of meiosis gene loci over synteny plots of the meiosis genes and 10 kb flanking regions upstream and downstream of the respective genes using EasyFig version 2.2.2 (64) between closely related species.

Furthermore, we compared possible relaxation of selection in the modeled meiosis genes of *Pseudozyma* (test set) to their homologs in sexually reproducing species (reference set) using RELAX (branch site model, version 3.1.1 (65)). Using gene-coding sequences and a species phylogeny, RELAX detects relaxation or intensification of selection between two different taxa by comparing the distribution of the ω values (dN/dS, ratio of non-synonymous substitutions and synonymous substitutions) estimated from a random effects branch site model (BS-REL) and classified in three categories (purifying, neutral and diversifying selection) from the test set in relation to a reference set. To reduce the complexity of the model, the values of ω in the test branches are set as ω_T_ = ω_R_^k^, with *k* the selection intensity parameter, so that *k*<1 indicates relaxed selection, *k*=1 no change in selection and *k*>1 intensification of selection in the test taxa. A likelihood ratio test is performed to assess whether the ω distributions of test and reference branches are significantly different, meaning that the test set would be under relaxed or intensified selection compared to the reference. We conducted this analysis only for the meiotic genes, because the different a-locus mating-type alleles have evolved before speciation, reducing the sample size to one specific mating-type and mating genes could be influenced by loss of the third allele in some species, while the b-locus genes show a too diverse allelicity in different species

## Declarations

### Ethics approval and consent to participate

Not applicable

### Consent for publication

Not applicable

### Availability of data and materials

The datasets supporting the conclusions of this article are available in the NCBI BioProject repository (ID: PRJNA816553; https://www.ncbi.nlm.nih.gov/bioproject/PRJNA816553 including BioSample SAMN26681043 to SAMN26681053) and in the figshare repository (doi: <u>10.6084/m9.figshare.19728994</u> and <u>10.6084/m9.figshare.20147135</u>).

Annotated gene sequences of Ustilaginales species are available in the DDBJ/ENA/GenBank databases under the accession numbers OP433763-OP434048 and the Third Party Annotation Section of the DDBJ/ENA/GenBank databases under the accession numbers TPA: BK062323-BK062532. Analyzed data that were not generated in this study are available at the accession numbers given in the manuscript.

### Author contribution

LS and DB conceived the study. TB supplied strains for sequencing. QW and FL conducted the cultivation, DNA isolation, and sequencing and assembled the genomes. LS reconstructed the phylogenetic tree and created manual and protein domain annotations and syntenies of meiosis genes. MG and LS annotated the mating-type loci and performed the assignment of mating-types. MD performed the selection analysis. LS wrote the manuscript. DB, MD, MG, and TB critically reviewed the manuscript. All authors read and approved the final manuscript.

## Acknowledgements

We acknowledge Magnus Wolf for providing assistance in the calculation of phylogenomic trees.

## Additional Files

Supplementary Figure 1: Phylogenetic tree of translated PRA gene sequences.

Supplementary Figure 2: Phylogenetic tree of translated MFa gene sequences.

Supplementary Figure 3: Synteny of core meiosis gene loci.

Supplementary Table 1: Meiosis and mating gene functional domains.

Supplementary Table 2: Telomere annotation.

Supplementary Table 3: Reference genes for mating gene annotation.

