## Supplementary Figure 1 for "Comparative genomics of smut fungi suggest the ability of meiosis and mating in asexual species of the genus *Pseudozyma* (Ustilaginales)"

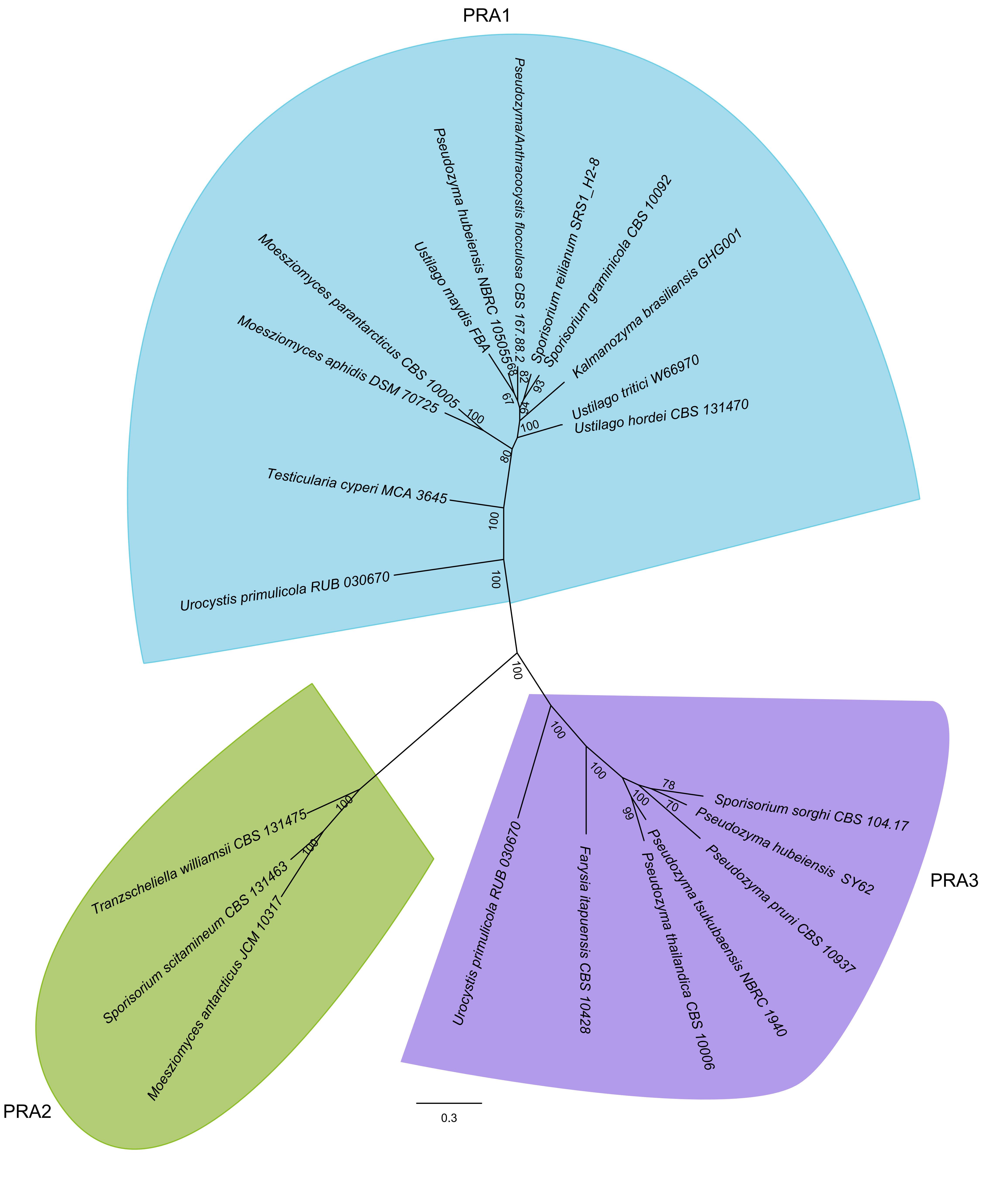


**Supplementary Figure 1:**
**Phylogenetic tree of translated PRA gene sequences.** The tree shows three clear clades of receptor gene alleles (one allele per species), allowing the annotation of mating-types for each strain.
