## Supplementary Figure 2 for "Comparative genomics of smut fungi suggest the ability of meiosis and mating in asexual species of the genus *Pseudozyma* (Ustilaginales)"

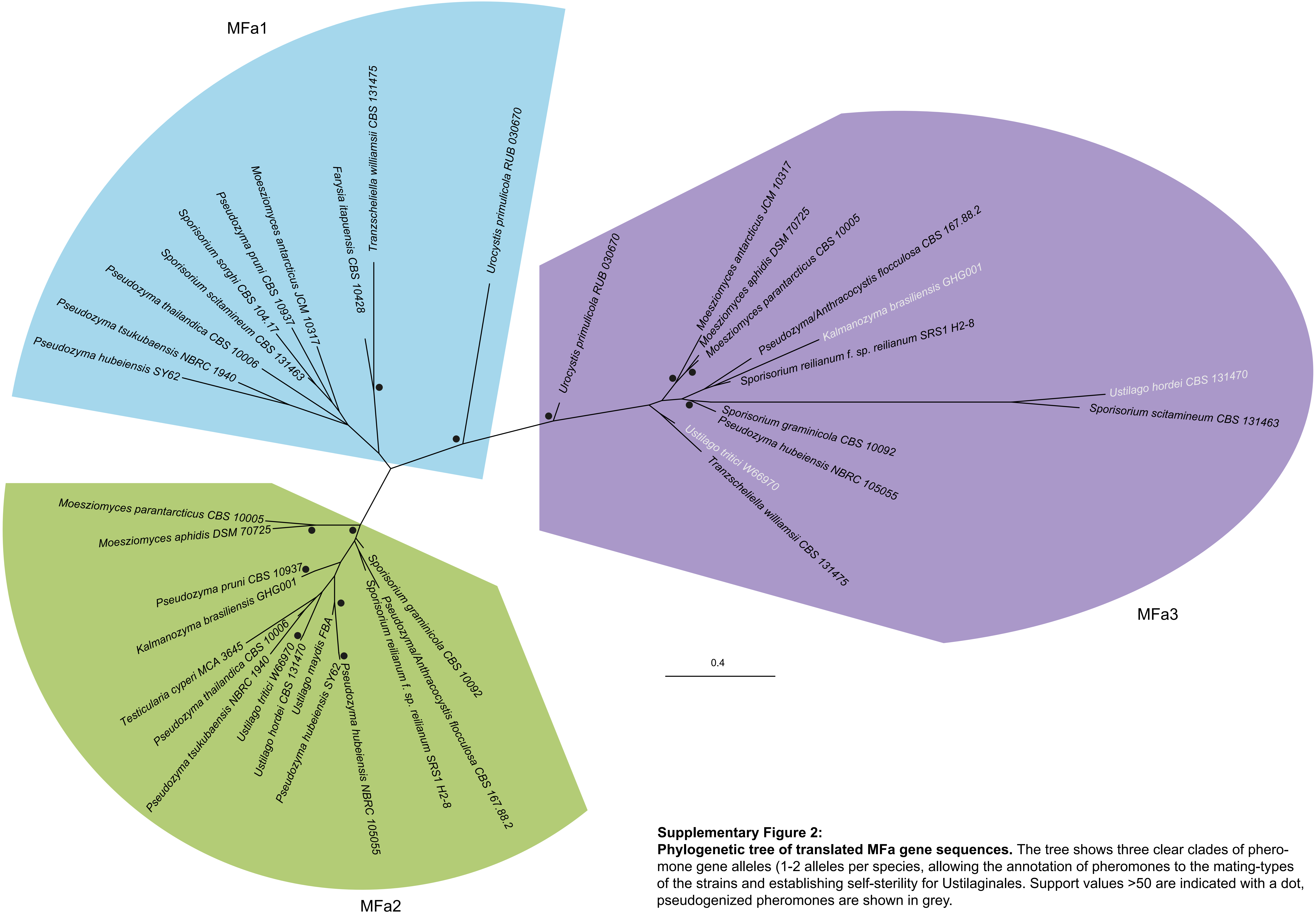


**Supplementary Figure 2:**
**Phylogenetic tree of translated MFa gene sequences.** The tree shows three clear clades of pheromone gene alleles (1-2 alleles per species, allowing the annotation of pheromones to the mating-types of the strains and establishing self-sterility for Ustilaginales. Support values >50 are indicated with a dot, pseudogenized pheromones are shown in grey.
