## Supplementary Figure 3 for "Comparative genomics of smut fungi suggest the ability of meiosis and mating in asexual species of the genus *Pseudozyma* (Ustilaginales)"

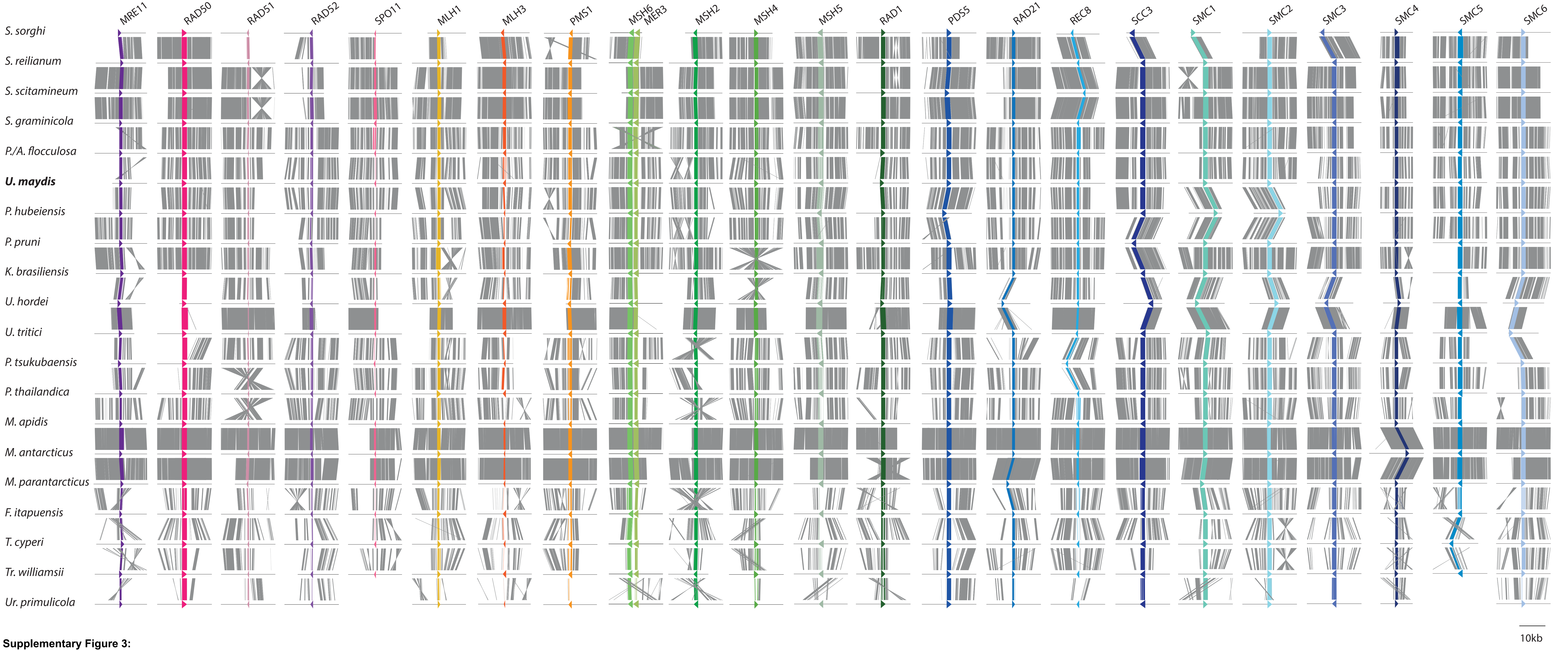


**Supplementary Figure 3:**
**Synteny of core meiosis gene loci.** Annotated genes and flanking regions (20 kb) remain mostly syntenic in relatively closely related species. More distantly related species like *Tr. williamsii*, *T. cyperi*, and the outgroup *Ur. primulicola* show less synteny. Rearrangements in the genomes of *Pseudozyma* compared to the sexual species cannot be noted. Species are sorted according to relatedness in the phylogenomic tree.
